# SLC25A46 is in contact with lysosomes and plays a role in mitochondrial cholesterol homeostasis

**DOI:** 10.1101/2024.04.01.587618

**Authors:** Jana Schuettpelz, Kathleen Watt, Hana Antonicka, Alexandre Janer, Ola Larsson, Eric A. Shoubridge

## Abstract

Mitochondrial morphology reflects the dynamic equilibrium between fusion and fission events, controlled by cellular signaling. A cytoprotective response known as stress-induced mitochondrial hyperfusion (SIMH) is triggered by nutrient starvation and we show that the outer mitochondrial membrane protein SLC25A46 is required for this response. To unravel the cellular mechanisms involved, we conducted transcriptomic analysis on control human fibroblasts and SLC25A46 knock-out cells. Our analysis revealed a remarkable divergence in the transcriptional profile of proteins associated with lysosomal function and cholesterol binding and synthesis. Further investigations using live-cell imaging validated the presence of SLC25A46 at the majority of mitochondria-lysosome contact sites. Since mitochondria-lysosome contacts are linked to cholesterol transport, we investigated the involvement of SLC25A46 in cholesterol trafficking. The SLC25A46 knock-out cell line exhibited a decrease in mitochondrial cholesterol content and distinct alterations were observed in the pattern of cholesterol trafficking compared to control. Cholesterol supplementation in the SLC25A46 knock-out cell line rescued the mitochondrial fragmentation phenotype and restored the SIMH response, suggesting a role for SLC25A46 in maintaining mitochondrial cholesterol homeostasis.

**Summary blurb:** The mitochondrial outer membrane protein SLC25A46 is required for SIMH triggered by nutrient starvation, localizes to lysosome contact sites and is involved in mitochondrial cholesterol homeostasis

## INTRODUCTION

Mitochondria form dynamic networks, balanced by fusion and fission events which respond to mitochondrial and cellular signalling (Shutt and McBride 2013). Mitochondrial division is essential for the elimination of damaged mitochondria (Twig et al. 2008; Youle and Narendra 2011), for organelle inheritance during proliferation (Twig et al. 2008), and the distribution of mitochondria within the cell (Mishra and Chan 2014). Fusion is thought to be important for mixing mitochondrial content (Chen, Chomyn, and Chan 2005), and is linked to an increased cellular energy production through oxidative phosphorylation (Yao et al. 2019) and mitochondrial import of fatty acids (Cohen, Rambold, and Lippincott-Schwartz 2018).

During mitochondrial fission cytosolic DRP1 binds to receptors (FIS1, MiD49, MiD51, MFF) on the outer mitochondrial membrane leading to the formation of oligomeric helices and subsequent mitochondrial constriction (Palmer et al. 2013). The fission site is marked by ER tubules, which may actively participate in this process (Friedman et al. 2011). Fusion on the other hand is dependent on the dynamin-like GTPases MFN1 and MFN2 in the outer membrane and OPA1 in the inner membrane (Cipolat et al. 2004). SLC25A46 is present at essentially all fission and fusion sites (Schuettpelz et al. 2023). Mitochondrial fragmentation and hyperfusion serve as mechanisms of quality control and survival responding to different cellular stresses (Shutt and McBride 2013) and has been observed in pathogenic infections (Stavru and Cossart 2011; Yang et al. 2020), cell death (Arnoult et al. 2005) and nutrient excess (Molina et al. 2009). Stress-induced mitochondrial hyperfusion (SIMH) is associated with ER stress (Lebeau et al. 2018), inhibition of translation (Tondera et al. 2009) and nutrient starvation (Gomes, Di Benedetto, and Scorrano 2011; Rambold et al. 2011). SIMH requires MFN1, OPA1 and stomatin-like protein 2 (SLP2) (Tondera et al. 2009); however, upstream regulatory mechanisms have not been identified.

While mitochondrial fusion and fission are coordinated at ER membrane contact sites (Abrisch et al. 2020), lysosomes have been described to play a role in mitochondrial fission as well (Wong, Ysselstein, and Krainc 2018). Mitochondria-lysosome tethering occurs following RAB7 GTP hydrolysis upon recruitment of TBC1D15 to the mitochondrial FIS1 receptor (Peralta, Martin, and Edinger 2010; Wong, Ysselstein, and Krainc 2018). The crosstalk between these two organelles includes the transfer of calcium, iron and cholesterol (Cisneros et al. 2022).

While the interactions between lysosomes and mitochondria are of great interest in cell biology, our current understanding of this inter-organelle contact site remains limited.

Autosomal-dominant mutations in *RAB7* lead to Charcot-Marie-Tooth (CMT) type 2B (Meggouh et al. 2006) while mutations of *MFN2* cause CMT type 2A. SLC25A46 has also been associated with CMT type 2 and a variety of other neurological disorders, such as optic atrophy and cerebellar atrophy (Abrams et al. 2018; Abrams et al. 2015; Braunisch et al. 2018; Buglo et al. 2020; Charlesworth et al. 2016; Hammer et al. 2017; Kodal et al. 2022). SLC25A46 is a mitochondrial outer membrane protein (Abrams et al. 2015; Janer et al. 2016) and plays an ill-defined role in mitochondrial fusion and fission, interacting with the fusion associated proteins MFN2 and OPA1, while the knock-out of SLC25A46 leads to altered oligomerization states of MFN2 and OPA1 (Schuettpelz et al. 2023).

We previously showed that the complete loss of SLC25A46 leads to mitochondrial fragmentation, while the expression of pathogenic variants leads to mitochondrial hyperfusion (Schuettpelz et al., 2023). Mitochondrial fusion has also been observed in patient cell lines with biallelic missense mutations (Abrams et al. 2015; Janer et al. 2016; Steffen et al. 2017) and following siRNA-mediated knock-down (Janer et al. 2016).

In this study we used transcriptomics, live-cell imaging, and biochemical analyses to further investigate the function of SLC25A46. We show that SLC25A46 is necessary for starvation-induced hyperfusion and that SLC25A46 is present at most mitochondria-lysosome contacts. Mitochondrial cholesterol is reduced and the mitochondrial fragmentation phenotype in SLC25A46 knock-out cells was rescued by cholesterol supplementation. These findings suggest that SLC25A46 is required for cholesterol homeostasis in mitochondria, the maintenance of which is required for mitochondrial fusion and fission.

## RESULTS

### SLC25A46 is required for SIMH during starvation

We previously reported that a complete loss of SLC25A46 in knock-out cell lines resulted in mitochondrial fragmentation, while re-expressing pathogenic variants in these cells led to a mitochondrial hyperfused network (Schuettpelz et al. 2023). Patients with biallelic missense mutations, which result in reduced expression of SLC25A46, also showed a fused mitochondrial network (Abrams et al. 2015; Janer et al. 2016; Steffen et al. 2017) which could be mimicked by acute siRNA-mediated knock down of SLC25A46. Thus, a complete lack of SLC25A46 leads to mitochondrial fragmentation while reduced expression leads to mitochondrial hyperfusion. Based on these data, we hypothesized that SLC25A46 might play a role in a protective SIMH response which can be induced by inhibition of translation or nutrient starvation (Gomes, Di Benedetto, and Scorrano 2011; Rambold et al. 2011; Tondera et al. 2009). In SLC25A46 knock-out cells treated with the translational inhibitor cycloheximide (CHX), mitochondria were predominantly hyperfused, similar to the control cell line, indicating that the fusion machinery is functional in SLC25A46 knock-out cells (Fig. 1a, b, c). However, SLC25A46 knock-out cells remained fragmented under nutrient starvation (HBSS), in contrast to the robust mitochondrial hyperfusion observed in the control cell line (Fig. 1a, b, c). To determine whether the mTOR pathway might be affected in the loss of SLC25A46 cells, we treated the cells with Ink128, an mTOR inhibitor known to effectively target both mTORC1 and mTORC2 complexes (Hsieh et al. 2012). To confirm the inhibition of the mTOR pathway, we analyzed the phosphorylation profile of 4E-BP, a direct target of the mTOR pathway. Immunoblot analysis showed a shift in the migration of 4E-BP on denaturing gels, indicating reduced phosphorylation by mTOR, confirming that Ink128 inhibits mTOR. (Fig. 1d). As expected, the control cells showed a mitochondrial hyperfusion phenotype after treatment with Ink128, an effect that was muted in the SLC25A46 knock-out cells suggesting that the distinct response to nutrient starvation in the SLC25A46 knock-out cell is not solely attributable to alterations in the mTOR pathway.

**Figure 1:**
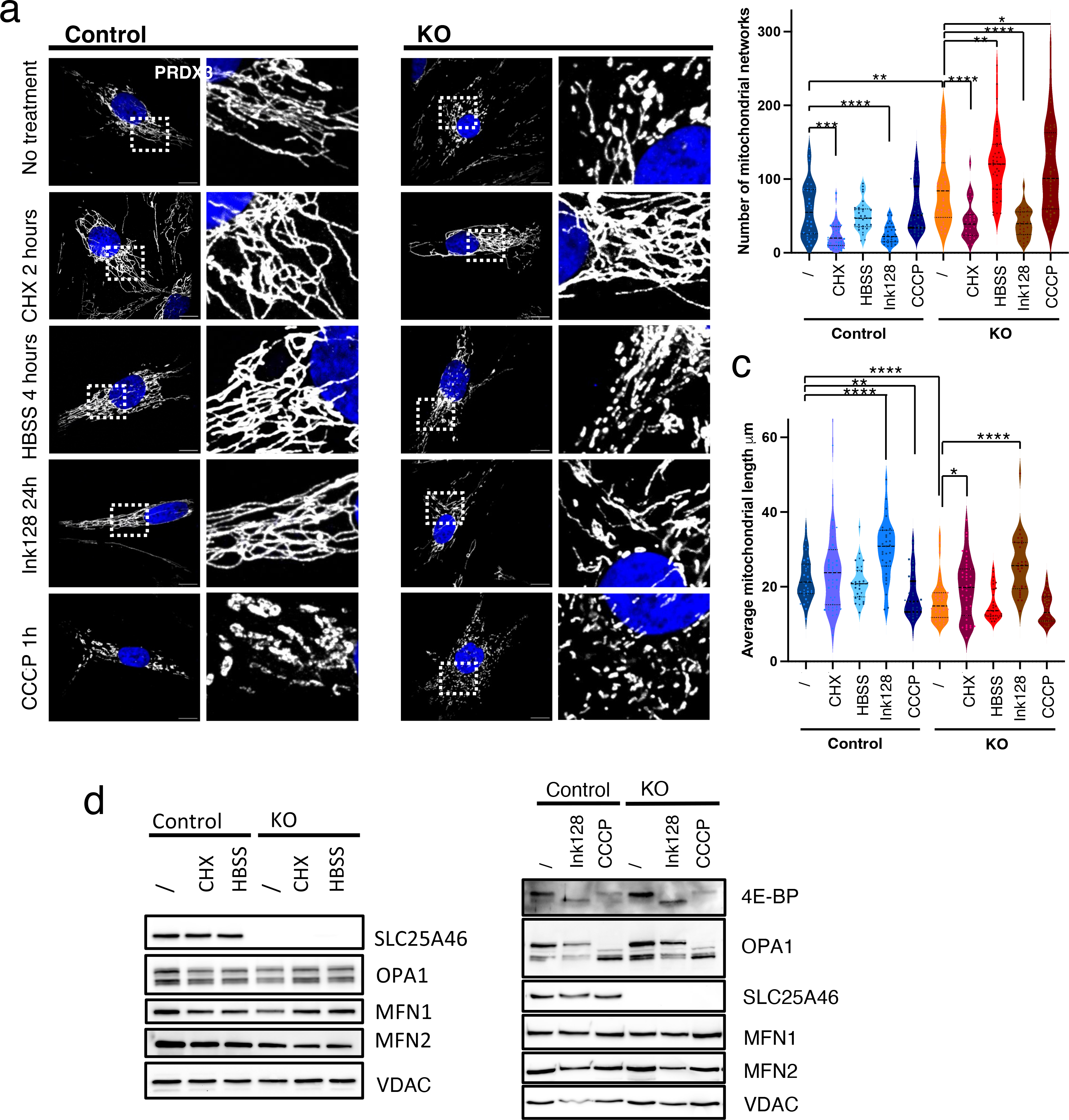
SLC25A46 is required for starvation-induced mitochondrial hyperfusion . The mitochondrial morphology of control fibroblasts and the SLC25A46 knock-out cell line was analyzed by immunofluorescence after treatment with cycloheximide (CHX) for 2 hours, with HBSS for 4 hours, with the mTOR inhibitor Ink128 for 24 h or CCCP for 1 hour. (a) Representative images of fibroblasts decorated with anti-PRDX3 (mitochondria) in white and DAPI (nuclei) in blue. A zoomed image of the indicated area is shown on the right. Scale bars: 10 µm. Images were analyzed using MitoMAPR (Zhang et al. 2019) to describe number of mitochondrial networks (b) and the average length of mitochondrial networks (c) (N=3, n=10) per condition shown as violin plots. (d) SDS-PAGE analysis of steady-state levels of indicated proteins in control fibroblasts and SLC25A46 knock-out cells after the indicated treatments.

To further investigate if stress-induced fragmentation is altered in SLC25A46 knock-out cells, we analyzed mitochondrial morphology following treatment with the protonophore carbonylcyanide m-chlorophenylhydrazone (CCCP). While mitochondria in both control and knockout exhibited mitochondrial fragmentation, the mitochondrial morphology was different:

SLC25A46 knock-out cells displayed smaller and more compact mitochondria, contrasting the larger and bulkier mitochondrial morphology observed in the control cells (Fig. 1a). Upon treatment with CCCP, OMA1 activity is enhanced resulting in proteolytic processing of OPA1 (Baker et al. 2014). The longer forms of OPA1 were degraded in the control cells as well as in the SLC25A46 knock-out cells (Fig. 1d). This finding implies that the stress response induced by CCCP may contribute to the observed differential mitochondrial morphology rather than affecting the processing of OPA1. OPA1 expression was not influenced by any other treatments, nor was the abundance of MFN1 and MFN2 (Fig. 1d). These data show that SLC25A46 is involved in mechanisms which lead to starvation-induced hyperfusion in an mTOR-independent manner.

### SLC25A46 knock-out cells have a different mRNA expression of lysosomal proteins

To investigate transcriptional alterations in the SLC25A46 knock-out cells, we determined the steady-state mRNA levels by RNA sequencing. To ensure consistency and avoid cell-specific differences, we also generated a SLC25A46 knock-out in HeLa cells by using the same gRNAs for CRISPR/Cas9 which we had used for the knock-out in fibroblasts.

Sequencing followed by differential expression analysis revealed dramatic changes in the transcriptome of both fibroblasts and HeLa cells with knock-out of SLC25A46 (Fig. 2a, Fig. S1a, b), with 4,287 and 3,986 differentially expressed genes (adjusted p-value < 0.1, Log2FC > |0.3|) between SLC25A46 knock-outs and controls in fibroblasts and HeLa cells, respectively. For further analyses, we focussed on the subset of 1,205 transcripts that exhibited significant changes in expression levels in both the SLC25A46 knock-out fibroblasts cell lines and the HeLa cell lines (Fig. 2b, Supplementary Table 1). This subset will be referred to hereafter as the SLC25A46-responsive genes.

**Figure 2:**
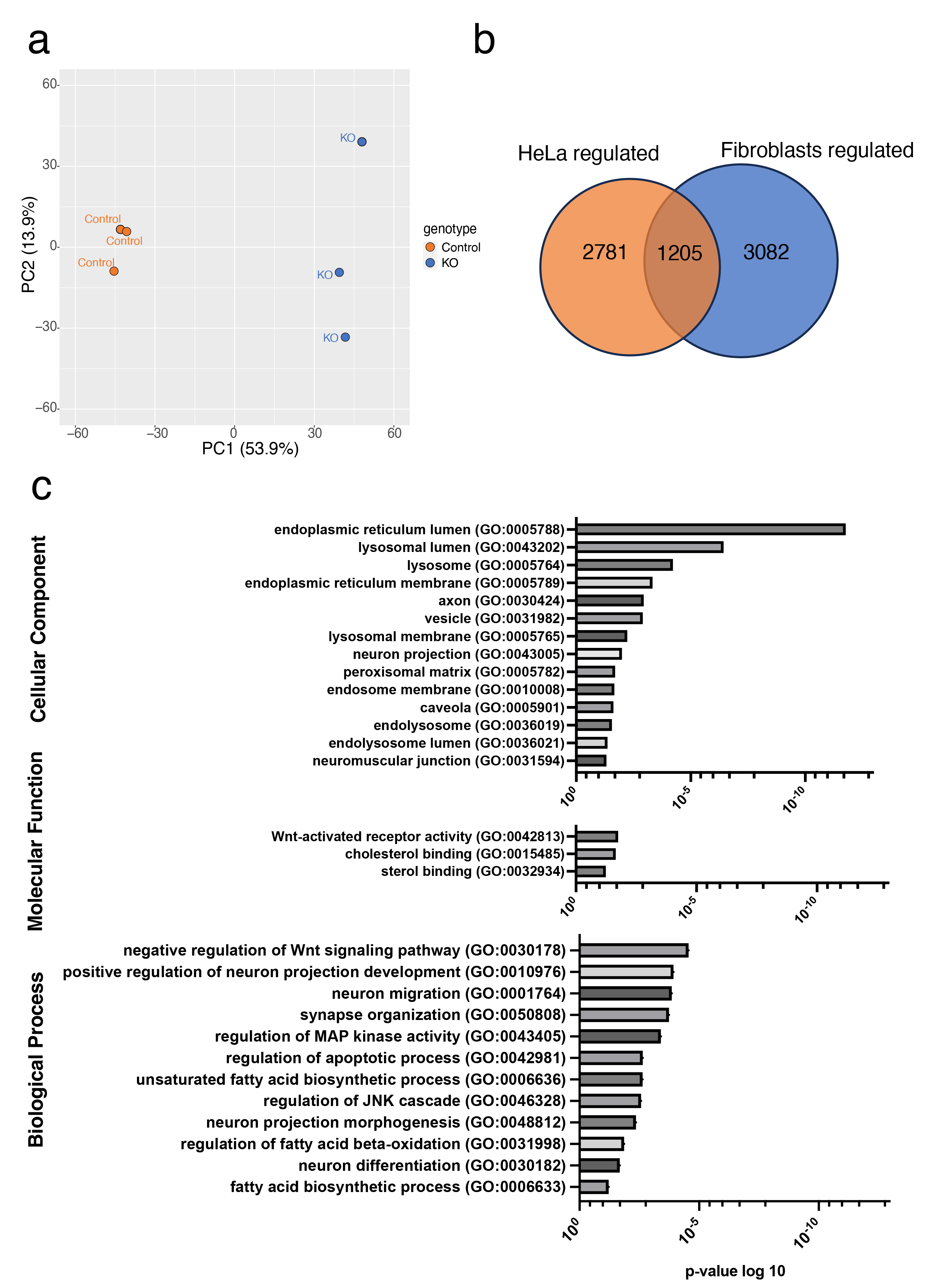
SLC25A46 knock-out cells have differential mRNA expression for genes encoding lysosomal proteins. Transcriptomic data was analysed for control and SLC25A46 knock-out cell lines in fibroblasts and HeLa cells. (a) Principle component analysis of gene expression quantified by RNA-seq in control and SLC25A46 knock-out fibroblast cells. (b) Only transcripts that were differently expressed in both cell lines (adjusted p-value < 0.1, Log2FC > |0.3|) were chosen to be analyzed (n = 1,790) (SLC25A46-responsive subset). (c) List of Gene Ontology (GO) terms with their p- value and the number of proteins of the group.

Gene Ontology enrichment analysis on the subset of SLC25A46-responsive genes revealed numerous transcripts encoding ER proteins with altered expression levels (Fig. 2c), so we next analyzed the ER in control and SLC25A46 knock-out fibroblast cell lines by immunofluorescence. Our observations did not reveal any obvious changes in ER structures in the absence of SLC25A46 (Fig. S2a). We found that transcripts coding for proteins expressed in vesicles (GO:00031982) and lysosomes (GO:0005764) (Fig. 2c, Fig. S2c, d) were highly dysregulated upon SLC25A46 knock-out, with many being upregulated in the SLC25A46 knock- out fibroblast cells lines in the subset of the SLC25A46-responsive genes. This corroborates our previous findings obtained by BioID proximity mapping showing that SLC25A46 is in proximity to proteins involved in vesicular transport (Schuettpelz et al. 2023). To further examine the lysosomal and mitochondrial dynamics we performed live-cell imaging in SLC25A46 knock-out cells expressing SLC25A46 with a C-terminal GFP-tag, focusing on dynamic interactions between mitochondria and lysosomes in SLC25A46 knock-out cells rescued with SLC25A46 tagged with GFP.

### SLC25A46 is in contact with lysosomes

Previous studies have demonstrated that approximately 15% of lysosomes in HeLa cells are in contact with mitochondria at any given time (Wong, Ysselstein, and Krainc 2018). We visualized mitochondria with MitoTracker®, lysosomes with LysoTracker® and SLC25A46 was followed using the GFP tag in fibroblasts (Fig. 3a). SLC25A46 was present at most mitochondria-lysosome contact sites (Fig. 3b). We detected a small but significant decrease in mitochondria-lysosome interactions in SLC25A46 knock-out fibroblasts compared to controls (Fig. 3c) indicating that while SLC25A46 might have a role in mitochondria-lysosome interactions, it is not required for those contacts. It was previously reported that lysosomes are present at most mitochondrial fission sites in HeLa cells (Wong, Ysselstein, and Krainc 2018); however, we were not able to confirm those observations in fibroblasts (Fig. 3d).

**Figure 3:**
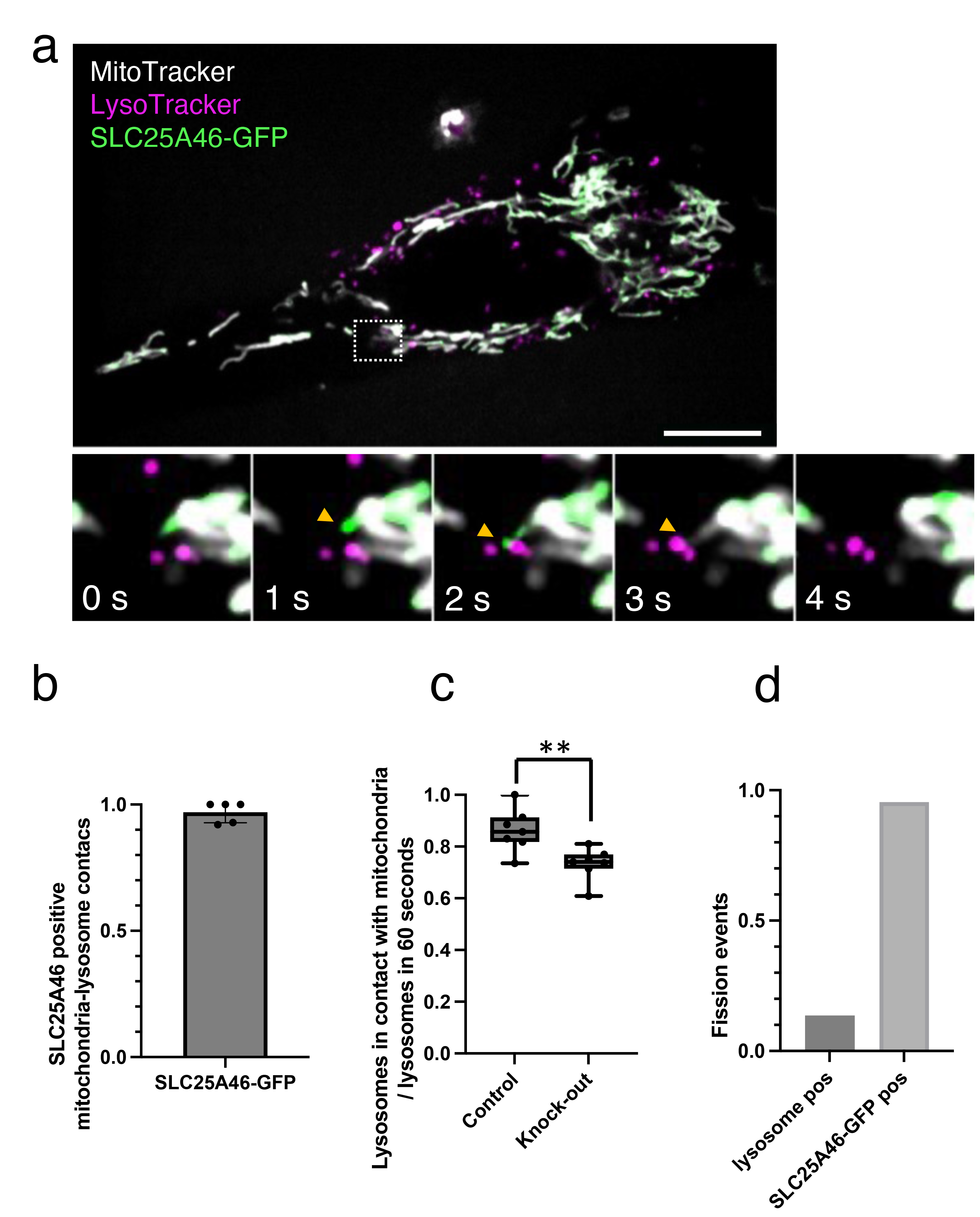
SLC25A46 is in contact with lysosomes. (a) Immunofluorescence analysis of SLC25A46 contacts with lysosomes in fibroblasts stably overexpressing SLC25A46-GFP (green). Mitochondria were stained with MitoTracker Deep Red (white) and lysosomes were stained with LysoTracker Red (magenta). Images were captured every 1 s for a period of 1 min. Time-lapse imaging of mitochondria-lysosome contact. Orange arrows indicate SLC25A46-GFP signal that contacts a lysosome. Scale bar: 10 μm. (b) Quantification of mitochondria-lysosome contacts positive for SLC25A46. Five cells were analyzed with a total of 94 mitochondria-lysosome contacts. (c) Quantification of the percentage of lysosomes in contact with mitochondria in control fibroblasts and in SLC25A46 knock-out cell line. Seven cells per condition were analyzed with a total of 230 lysosomes in control cells and 188 lysosomes in knock-out cells. (d) Quantification of mitochondrial fission sites with a lysosome present (lysosome pos) and positive for SLC25A46 (SLC25A46-GFP pos). Seven cells were analysed with 22 fission events. Data shown as mean+SEM.

### SLC25A46 influences mitochondrial cholesterol homeostasis

We previously showed a significantly different mitochondrial lipid content in the SLC25A46 knock-out cells compared to controls (Schuettpelz et al. 2023) which prompted us to examine our transcriptomic dataset for alterations in transcripts related to lipid homeostasis. Our analysis revealed a substantial number of differentially expressed SLC25A46-responsive transcripts encoded for proteins involved in fatty acid biosynthesis (GO:0006633), a GO-term including proteins involved in cholesterol synthesis as well (Fig. 2c, 4a). Several transcripts associated with cholesterol binding (GO:0015485) were upregulated (Fig. 2c, 4b).

Since our previous lipidomic analysis did not include the analysis of the mitochondrial cholesterol content, we measured the cholesterol levels in control cells, SLC25A46 knock-out cells and SLC25A46 knock-out cells expressing wild-type SLC25A46 (described in earlier studies (Schuettpelz et al. 2023)) (Fig. 4c, d). The cholesterol content of whole cells was not significantly changed (Fig. 4c); however, the free cholesterol content in isolated mitochondria was significantly decreased in the absence of SLC25A46 and this was rescued by expressing the wild-type protein (Fig. 4d).

**Figure 4.**
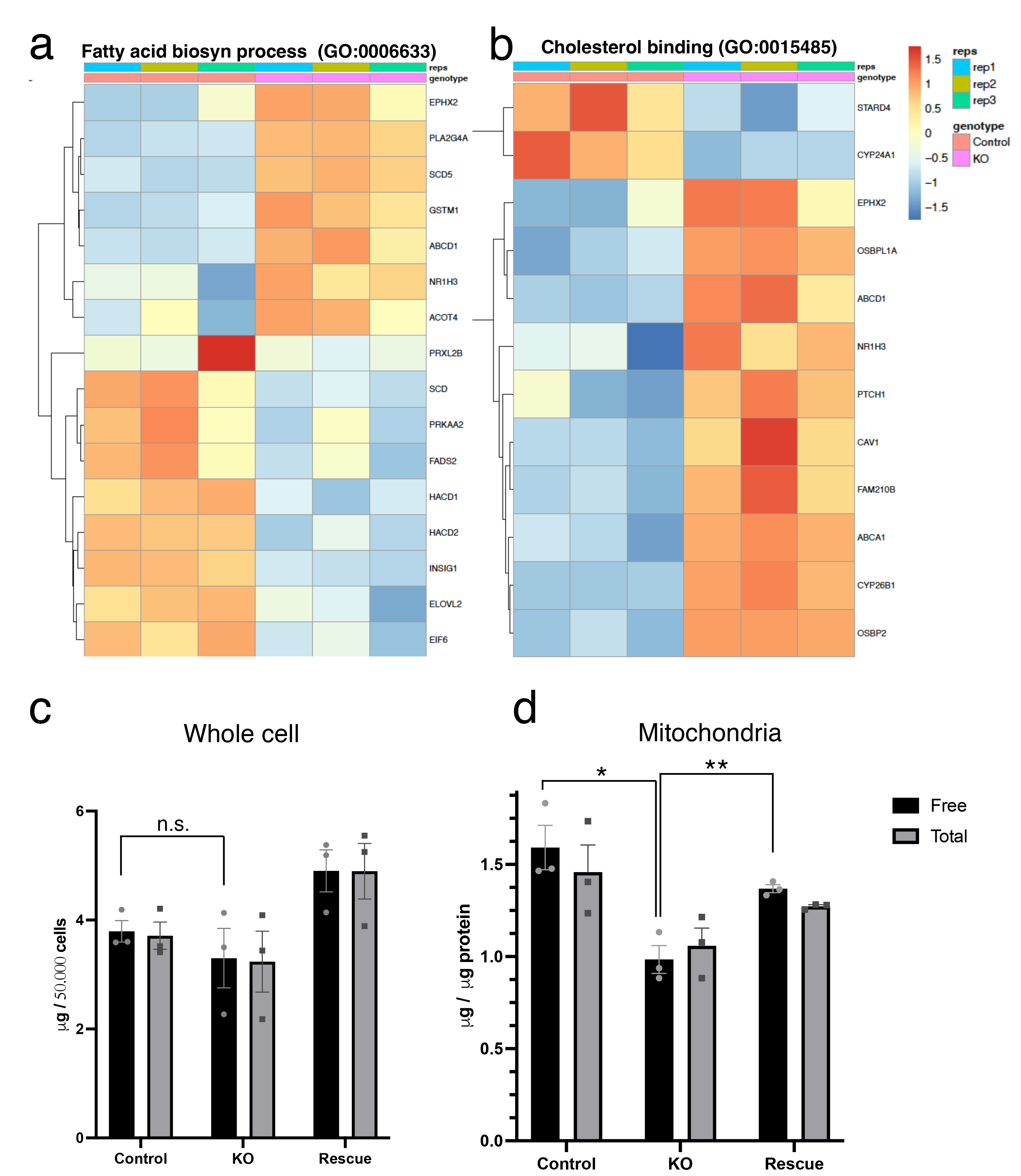
SLC25A46 influences mitochondrial cholesterol homeostasis. Heatmap of differently expressed (adjusted p-value < 0.1, Log2FC > |0.3|) SLC25A46- responsive transcripts in control and SLC25A46 knock-out fibroblasts coding for proteins involved in fatty acid biosynthesis (GO:0006633) (a) and cholesterol binding (GO:0015485) (b). Log2 fold changes are centered and represented as z-scores. Total cholesterol and non- esterified (“free”) cholesterol in whole cells (c) and in sucrose-gradient isolated mitochondria (d) from control fibroblasts, SLC25A46 knock-out cells (KO) and SLC25A46 knock-out cells expressing the wildtype SLC25A46 protein (Rescue) was measured with a Cholesterol/Cholesteryl Ester Assay Kit (Abcam, ab65359).

### Cholesterol trafficking is dependent on SLC25A46

To test whether SLC25A46 is involved in cholesterol trafficking we analysed the kinetics of cholesterol import and trafficking using the fluorescent cholesterol probe TopFluor® in control and SLC25A46 knock-out cells (Fig. 5a, b). After 30 min incubation the cholesterol probe appeared in mitochondria, plasma membrane and some lysosomal structures then migrated to lysosomal structures after 1 hour (Fig. 5a, Fig. S2b). While in the control cells the fluorescent probe stayed in lysosomal structures, in the knock-out cells it appeared in mitochondria and plasma membrane again after 1.5 hours. The signal only appeared back in lysosomes after 2.5 hours indicating that the cholesterol trafficking is altered in absence of SLC25A46 (Fig. 5a, b, Fig. S2b).

**Figure 5:**
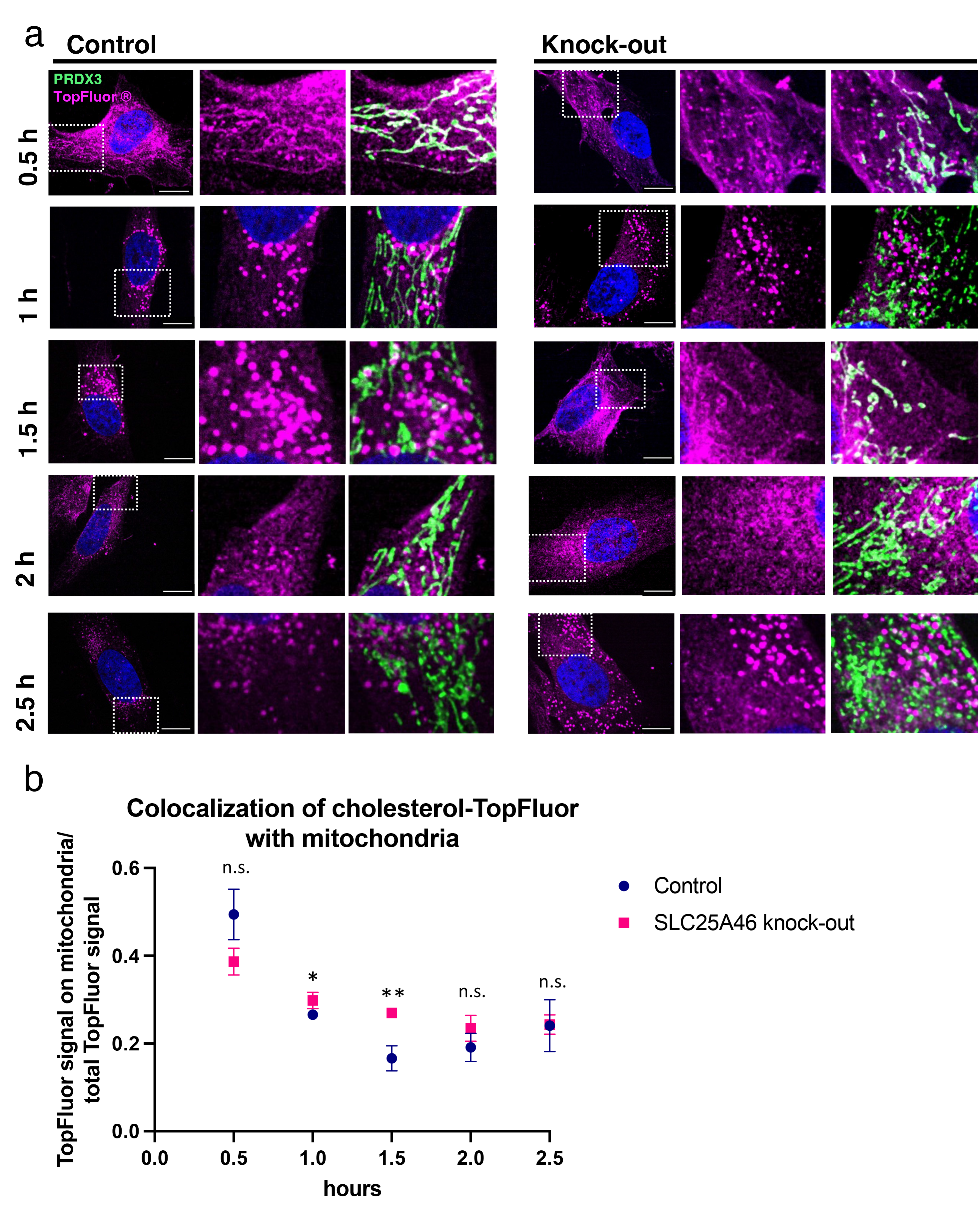
Cholesterol trafficking is dependent on SLC25A46. Immunofluorescence analysis of cholesterol trafficking in control human fibroblasts and SLC25A46 knock-out cells. Cells were pulsed for 2 min with the fluorescent cholesterol probe TopFluor® (in magenta) and fixed after the indicated time to track the probe. PRDX3 was used as a mitochondrial marker (green). A zoomed image of the boxed area is shown on the right. Scale bars: 10 μm. (b) Quantification of colocalizing TopFluor signal with mitochondria divided by the total TopFluor signal in the cell at indicated time points. n=10 per condition.

### Cholesterol can rescue the mitochondrial fragmentation phenotype of the SLC25A46 knock-out cell line

The above data suggested a possible role of SLC25A46 in cellular cholesterol trafficking to mitochondria and as the SLC25A46 knock-out cells show a fragmented mitochondrial phenotype we analyzed mitochondrial morphology in control and SLC25A46 knock-out cells after treating the cells with 20 μM cholesterol for 4 hours. Strikingly, the mitochondrial morphology of the SLC25A46 knock-out cell line was rescued while in the control cell line the morphology became less hyperfused as the average length was decreased (Fig. 6a, b, c).

**Figure 6:**
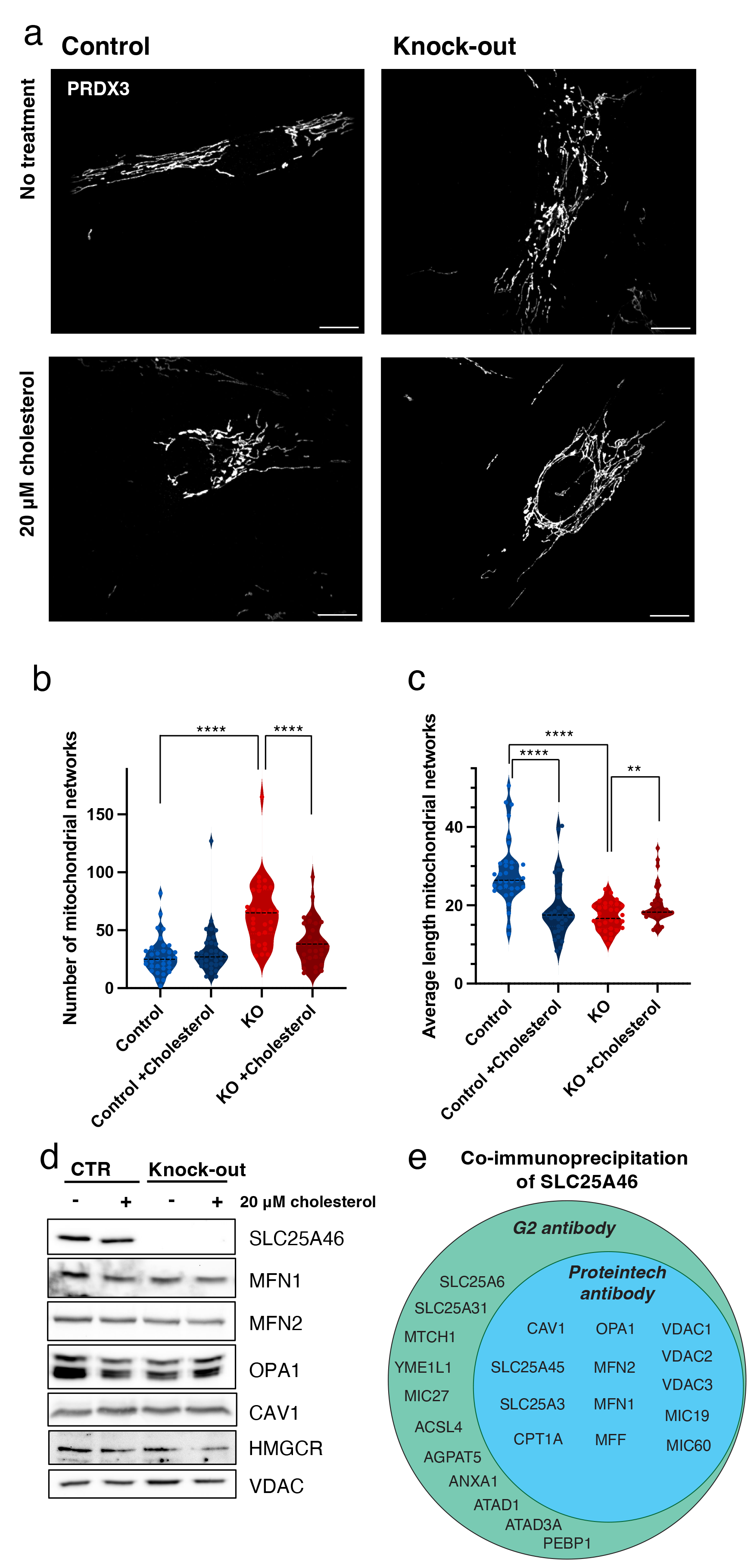
Cholesterol rescues the mitochondrial fragmentation phenotype of the SLC25A46 knock-out cell line and interaction of SLC25A46 with Cav1. The mitochondrial morphology of control and SLC25A46 human fibroblasts was analyzed after treating the cells with 20 μM cholesterol for 4h. Representative immunofluorescent images with the mitochondrial marker PRDX3 (white) (a). Scale bars: 10 μm. Images were analyzed using MitoMAPR (Zhang et al. 2019) across max projections keeping all parameters constant (N=3, n=30) to describe number of mitochondrial networks (b) and the average length of mitochondrial networks (c) per condition shown as violin plots. SDS-PAGE analysis of control and SLC25A46 knock-out cell lines (d). VDAC was used as a loading control. (e) Venn diagram of proteins co- immunoprecipitated with SLC25A46. Two different antibodies (G2: Santa Cruz; G2-SC-515823 and Proteintech: 12277-1-AP) against the endogenous SLC25A46 protein we used for immunoprecipitation of crude mitochondria isolated from control fibroblasts. The co- immunoprecipitated proteins were identified by mass spectrometry.

We next examined the effects of cholesterol treatment on key components of cholesterol homeostasis by performing SDS-PAGE on the cell lysates of control cells and the SLC25A46 knock-out cell lines. Since the mRNA of Caveolin-1 (Cav1) was upregulated in the SLC25A46 knock-out cell line (Fig. 4b) we analyzed the protein abundance of this protein. Cav1 is an integral membrane protein which is required to generate membrane curvatures such as caveolae and plays an essential role in the regulation of cellular cholesterol metabolism (Cheng and Nichols 2016); however, the protein levels of Cav1 were not altered in the SLC25A46 knock-out cell-line (Fig. 6d, Fig. S1b). The rate-limiting enzyme HMGCR (HMG-CoA reductase) in cellular cholesterol synthesis in the ER (Arenas, Garcia-Ruiz, and Fernandez-Checa 2017) was also not changed in the SLC25A46 knock-out cell line (Fig. 6d, Fig. S1b), suggesting that the steady-state levels of proteins involved in cholesterol synthesis and the caveolae are not dependent on the presence of SLC25A46 abundance or affected by short-term supplementation of cholesterol.

Furthermore, through immunoprecipitation experiments, we confirmed an interaction between SLC25A46 and Cav1. In our previous studies using mass spectrometry and immunoblot analysis we demonstrated that SLC25A46 immunoprecipitates proteins of the fusion machinery and components of the MICOS complex (Janer et al. 2016; Schuettpelz et al. 2023). In these two experiments we used two different antibodies (G2 and Proteintech), both indicating an interaction of SLC25A46 with Cav1 (Fig. 6e, Supplementary Table 2). Additionally, when we immunoprecipitated Cav1, we also detect SLC25A46, albeit with a low score (Supplementary Table 2). These data suggest a possible association between these two proteins.

### Transcriptomics show an altered cellular response during nutrient starvation and cholesterol can induce a SIMH response in the SLC25A46 knock-out cells

Our results showed a different response to SIMH in SLC25A46 knock-out cells compared to our control cell line, especially upon nutrient starvation (HBSS) (Fig. 1a, b, c, Fig. S1c). To interrogate the cellular signalling of these different responses, we analyzed the transcriptome of control cells and the SLC25A46 knock-out cell line upon treatment with HBSS. Both control and SLC25A46 cell lines treated with HBSS exhibited robust responses as evidenced by significant changes in expression of thousands of genes (n = 5,116 and n = 5,588 respectively, at adjusted p-value < 0.1 and Log2FC > |0.3|). We also identified a large subset of genes where HBSS-induced changes in expression were dependent on SLC25A46 (n = 1,612, at adjusted p-value < 0.1 and Log2FC > |0.3|) (Fig 7a). Among this subset of genes for which stress-induced changes in expression were dependent on SLC25A46 were transcripts encoding proteins in the GO-terms of lysosomes (GO:0005764), fatty acid biosynthesis (GO:0006633) and cholesterol binding (GO:0015485) (Fig. 7b, c, d, Supplementary Table 1). Furthermore, we also noted differential regulation upon HBSS treatment between SLC25A46 knock-out and control cell lines for genes involved in NF-κB signalling, response to ER-stress, MAPK-cascade and JNK cascade (Fig. 7e, Fig. S3), indicating that the absence of SLC25A46 leads to altered cellular responses to stress during starvation compared to control cells.

**Figure 7:**
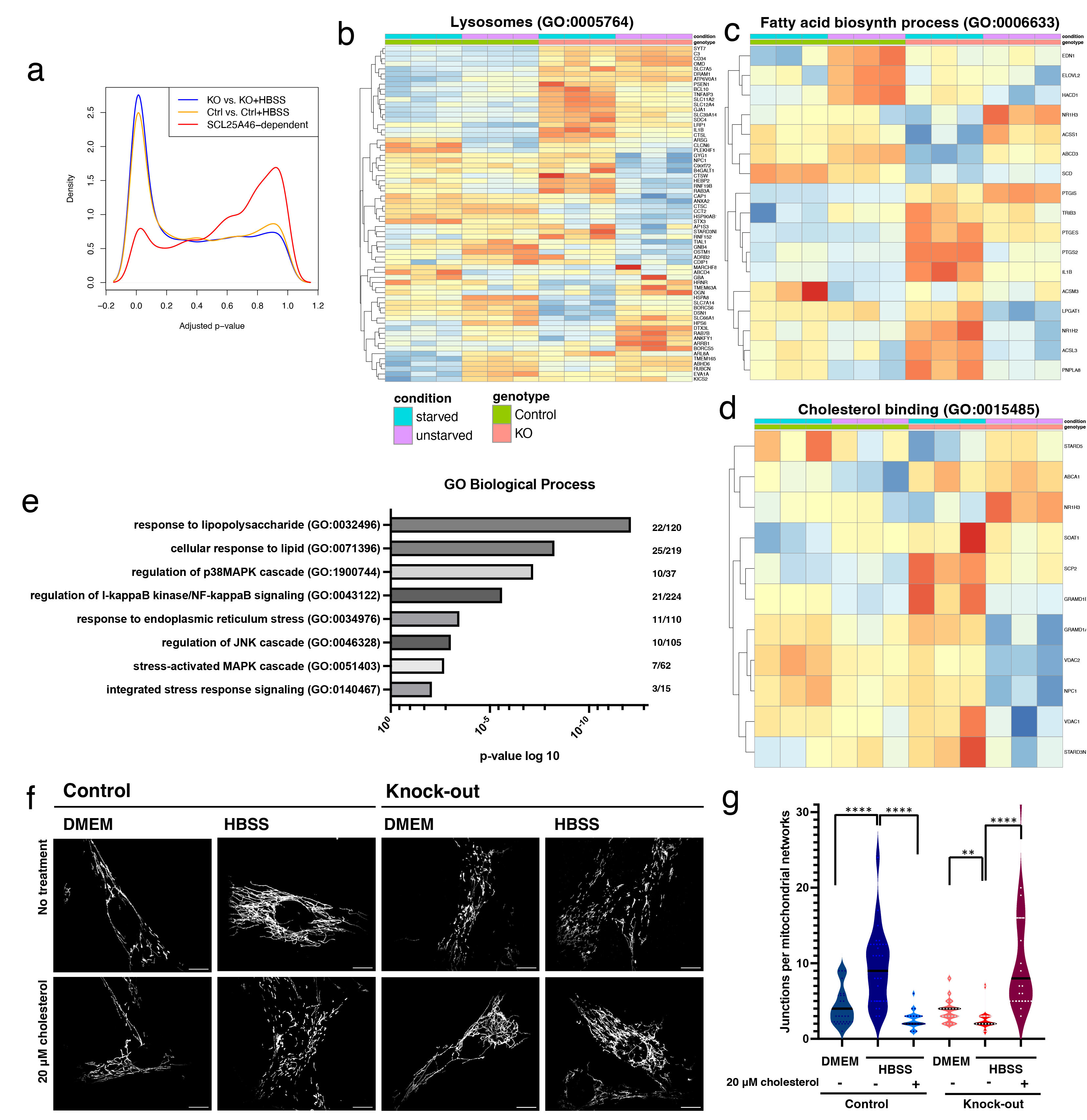
An altered cellular response during nutrient starvation in the SLC25A46 knock- out cells. (a) Kernel density estimation adjusted p-value distributions for differentially expression genes in control and SLC25A46 KO cells upon treatment with HBSS, as well as genes where the differential expression upon HBSS treatment was dependent on SLC25A46 expression. A higher density of low adjusted p-values is indicative of enrichment in changes in gene expression. Heatmaps of changes in levels of transcripts of lysosomal proteins (GO:0005764) (b), proteins involved in fatty acid biosynthesis (GO:0006633) (c) and in cholesterol binding (GO:0015485) (d) in control and SLC25A46 knock-out fibroblasts in normal conditions and during nutrient starvation. Genes displayed are those where HBSS-induced changes in expression were dependent on SLC25A46 in the DESeq2 model with the design “expression ∼ genotype + condition + genotype x condition” (adjusted p-value < 0.1 and Log2FC > |0.3| for the “genotype x condition” term). Log2 fold changes are centered and represented as z-scores. (e) List of Gene Ontology (GO) terms with their p-value and the number of proteins in the group. (f) Representative images of the mitochondrial morphology of control and SLC25A46 human fibroblasts after 4 hours of incubation in normal medium (DMEM) or under nutrient starvation (HBSS), either without supplementation (no treatment) or with supplementation of 20 μM cholesterol for 4h. Mitochondrial marker PRDX3 (white). Scale bars: 10 μm. (g) Images were analyzed using MitoMAPR (Zhang et al. 2019) on ROIs keeping all parameters constant to quantify number of junction per mitochondrial networks (N=3, n=30).

To determine if cholesterol supplementation would rescue the SIMH response in the SLC25A46 knock-out cell line, we analysed mitochondrial morphology by immunofluorescence confocal imaging. While nutrient starvation (HBSS) in the SLC25A46 knock-out cell lines led to mitochondrial fragmentation, the control cell line responded with mitochondrial elongation (Fig. 1 and Fig. 6). Addition of cholesterol to the HBSS medium in the SLC25A46 knock-out cell line resulted in an elongated mitochondrial morphology, similar to the control cell line undergoing nutrient starvation (in HBSS alone). On the other hand, supplementation of cholesterol in the control cell line resulted in mitochondrial fragmentation (Fig. 7f, g). This response could potentially be attributed to an excess accumulation of cholesterol, as previously described by Solsona-Vilarrasa et al. (Solsona-Vilarrasa et al. 2019). These data indicate that the function of SLC25A46 is linked to mitochondrial cholesterol homeostasis, which may be necessary for SIMH.

## DISCUSSION

Although SLC25A46 is present at essentially all mitochondrial fission and fusion sites, our previous studies (2023) showed that mitochondria fragment in SLC25A46 knockout cells, whereas hypomorphic loss of function missense mutations elicit mitochondrial hyperfusion.

These data demonstrate that SLC25A46 plays a nonessential role in the mitochondrial fission and fusion machinery per se. Further, mitochondria in SLC25A46 knock-out cells treated with cycloheximide were predominantly hyperfused, similar to controls indicating a functional fusion machinery. (Fig. 1a) (Schuettpelz et al. 2023). Nutrient starvation invariably promotes a protective starvation induced mitochondrial hyperfusion response in control cells (Gomes, Di Benedetto, and Scorrano 2011; Rambold et al. 2011); however, mitochondria in SLC25A46 knock-out cells remained fragmented under conditions of nutrient starvation (Fig. 1a) suggesting that SLC25A46 plays a selective role in the starvation-dependent SIMH pathway, possibly due to the lack of a specific component in mitochondria. The results of this study suggest that this component may be cholesterol as its supplementation successfully restored the impaired SIMH response observed in the SLC25A46 knock-out cell line (Fig. 7).

The enrichment of cholesterol in hepatocyte mitochondria was associated with a fragmented morphology, along with impaired assembly of the OXPHOS complexes and decreased respiration (Solsona-Vilarrasa et al. 2019). Similar to these findings, we observed a more fragmented morphology when we supplemented control fibroblast cells with cholesterol. On the contrary, when the SLC25A46 knock-out cell line was supplemented with cholesterol, we observed a more elongated mitochondrial network demonstrating that cholesterol supplementation rescues the fragmented mitochondrial morphology induced by the loss of SLC25A46.

We also show that SLC25A46 is present at most lysosome-mitochondria contact sites (Fig. 3) and our previous study has shown that SLC25A46 is present at 98% of all fusion and fission sites (Schuettpelz et al. 2023). Lysosomes have been reported to be present at ∼80% of fission events in HeLa cells (Wong, Ysselstein, and Krainc 2018); however, we were unable to reproduce this observation in fibroblasts (Fig. 3c). Consequently, we can confidently rule out the possibility that SLC25A46-lysosome interactions happen solely due to their simultaneous occurrence at mitochondrial fission/fusion events. Instead, SLC25A46 foci predominantly exhibited a distinct “kiss-and-run” interaction pattern, wherein a mitochondrial tip carrying a SLC25A46-GFP positive extrusion briefly interacted with a lysosome (exemplified in Fig. 3a).

We observed a reduction of approximately 10% in the number of mitochondria-lysosome contacts in SLC25A46 knock-out cells compared to controls (Fig. 3d) indicating that while SLC25A46 may not be indispensable for establishing direct mitochondria-lysosome contacts, its function might operate downstream of transport processes occurring between these organelles. Qiu et al. described an elongated mitochondrial phenotype in their SLC25A46 knock-out cell line and proposed that by inducing mitochondria-lysosome contacts, they could rescue the mitochondrial hyperfusion phenotype (Qiu et al. 2022) providing additional support for the involvement of SLC25A46 in modulating mitochondrial dynamics linked to mitochondria-lysosome contacts.

SLC25A46 knock-out cells showed a decrease in mitochondrial cholesterol content and the addition of exogenous cholesterol rescued the fragmentation phenotype (Fig. 4d, 6a, b, c), suggesting that SLC25A46 plays an important role in mitochondrial cholesterol homeostasis.

Duchesne et al. (2017) conducted biochemical blood analysis on SLC25A46 knock-out mice, revealing elevated cholesterol levels compared to control mice (5.85 ± 1.40 mmol/L vs. 3.25 ± 0.19) suggesting a disturbance in cellular uptake or metabolism of cholesterol in the absence of SLC25A46.

Cholesterol is essential for certain properties of the cell membrane, influencing the membrane fluidity, permeability and curvature. Furthermore, the level of cholesterol can affect protein functions and interactions. While cholesterol is predominantly found in the plasma membrane and Golgi, its *de novo* synthesis takes place in the ER by several enzymatic reactions. Despite the fact that the ER and mitochondrial membranes contain approximately 40 times less cholesterol than the plasma membrane, its scarcity does not diminish its importance (Martin, Kennedy, and Karten 2016). Mitochondria have the ability to acquire cholesterol from various subcellular membranes, resulting in a redundancy in the import pathways for mitochondrial cholesterol. The import of cholesterol to mitochondria can occur through carrier-mediated transport and movement across membrane contact sites (Elustondo, Martin, and Karten 2017).

Fluorescently tagged cholesterol tracking demonstrated that, different to the control cell line, cholesterol was observed to return from lysosomes to mitochondria within the timeframe of 1.5 to 2.5 hours in the absence of SLC25A46 (Fig. 5), suggesting that the absence of SLC25A46 may trigger an alternative pathway for cholesterol import. The additional cholesterol efflux back from lysosomes to mitochondria in the absence of SLC25A46 between 1.5 h and 2.5 h does not necessarily align with the general mitochondrial cholesterol decrease (Fig. 4).

The findings in SLC25A46 knockout cells differ from the mitochondrial cholesterol accumulation observed in the lysosomal storage disease Niemann-Pickmann Disease C (NPC) caused by defects in the lysosomal proteins NPC1 or NPC2 which are involved in cholesterol efflux from lysosomes. NPC is characterized by a mitochondrial cholesterol accumulation (Torres et al. 2017). This contrast suggests that the role of SLC25A46 in cholesterol trafficking may involve distinct mechanisms or pathways compared to those disrupted in NPC. The trafficking of fluorescent cholesterol probes has not been studied to any extent in different cell types (Kralova et al. 2018). The precise details of cholesterol trafficking between the ER, mitochondria, and lysosomes are largely unknown and involve redundant pathways with protein-mediated transfer, endosomal/lysosomal trafficking, and membrane contact sites (Elustondo, Martin, and Karten 2017). While it is evident that cholesterol cycling is affected in the absence of SLC25A46, the underlying mechanisms remain to be investigated.

CAV1, associated with caveolae and lipid droplets, plays a critical role in maintaining unesterified cholesterol levels (Cheng and Nichols 2016; Parton and Simons 2007). We found that CAV1 interacts with SLC25A46 (Figure 6e, Supplementary Table 1) and that its mRNA expression was increased in SLC25A46 knockout cells (Fig 4b and 6d). CAV1 is known to transport cholesterol from mitochondria to the plasma membrane as the knock-out of CAV1 leads to mitochondrial cholesterol accumulation and deficiency is the plasma membrane (Bosch et al. 2011). These observations suggest an intriguing pattern: while the levels of mitochondrial cholesterol are decreased, there is an increase in transcripts of proteins involved in cholesterol export from mitochondria. These findings implicate a potential problem in the upstream regulation of cholesterol homeostasis. Overall, our findings highlight the complexity of cholesterol trafficking and emphasize the potential role of SLC25A46 in modulating mitochondrial cholesterol dynamics through its interactions with lysosomes, influencing mitochondrial morphology.

## MATERIALS AND METHODS

### Cell culture

Fibroblasts were obtained from a cell bank located in the Montreal Children’s Hospital and the cell line used was from a female healthy subject, 58 years old. The cells were immortalized as described previously (Lochmuller, Johns, and Shoubridge 1999). The generation of CRISPR- Cas9 SLC25A46 knockout cell line, knock-out cell line over-expressing wild-type SLC25A46 (Rescue), as well as cell line over-expressing SLC25A46-GFP were described previously (Schuettpelz et al. 2023). The HeLa SLC25A46 knock-out cell line was generated with the same gRNA as the fibroblast cell line (Schuettpelz et al. 2023).

All fibroblast- and HeLa-derived cell lines were cultured in high-glucose DMEM (Wisent 319- 027-CL) supplemented with 10% fetal bovine serum, at 37°C in atmosphere of 5% CO2. All cell lines were regularly tested for mycoplasma contamination.

For SIMH experiments, cells were treated with 200 µM cycloheximide in complete medium or washed three times in HBSS (Gibco) to remove any residual medium, then kept in HBSS. Where indicated, cells were incubated with 100 nM CCCP for 1h or Ink128 (MLN0128, AdooQ) for 24 h.

For the cholesterol treatment, cells were incubated with DMEM or HBSS supplemented with 20 µM cholesterol (Sigma-Aldrich; C4951) for 4 h.

### Preparation of mitochondria enriched heavy membranes

Mitochondria enriched heavy membrane fractions were acquired from two 90% confluent 15 cm plates. The plates were washed twice with PBS, cells were collected and resuspended in HIM buffer (0.2M mannitol, 0.07M sucrose, 0.01M HEPES, 1mM EGTA, pH 7.5) and homogenized with ten passes of a pre-chilled, zero-clearance homogenizer (Kimble/Kontes). The post-nuclear supernatant was obtained by centrifugation of the samples twice for 10 min at 600g. The heavy membrane fraction was pelleted by centrifugation for 10 min at 10,000g and washed once in HIM buffer. The protein concentration was measured by Bradford assay.

### Denaturating, native and second dimension PAGE

BN-PAGE was used to separate individual protein complexes. Each sample was prepared from 100 μg mitochondria enriched heavy membrane fractions. Mitochondria were resuspended in 40 μl MB2 buffer (1.75M aminocaproic acid, 75 mM Bis-tris, 2mM EDTA) and solubilized with 4 μl 10% digitonin solution (4g digitonin/g protein) for 20 min on ice with subsequent centrifugation for 20 min at 20,000g. 15 μg of supernatants were separated on a 6–15% polyacrylamide gradient gel as previously described (Leary 2012), with Amersham HMW native marker kit (17044501; GE Healthcare) as a molecular weight marker, and blotted onto a polyvinylidene difluoride (PVDF) membrane. For SDS–PAGE, 15 μg of protein was run on polyacrylamide gels. For the second-dimension analysis, BN–PAGE/SDS–PAGE was carried out as detailed previously (Antonicka et al. 2003). Separated proteins were transferred to a nitrocellulose membrane and immunoblot analysis was performed with the indicated antibodies.

### Cholesterol measurement

For measurement of cholesterol in whole cell extract 10^6^ cells per condition were washed three times with cold PBS and harvested. The cells were lysed in 200 μL of a chloroform/isopropanol/NP-40 solution (7:11:0.1) in a micro-homogenizer with subsequent centrifugation for 10 min at 15,000g. The supernatant was vacuum dried and the dried lipids were dissolved in 200 μL of the assay buffer supplied by the Cholesterol/ Cholesteryl Ester Assay Kit (Abcam, ab65359). 5 μL of each sample was used for each reaction.

To obtain at least 100 μg of isolated mitochondria from fibroblasts, 4 x 15cm plates with a 90% confluency were grown for each sample. The plates were rinsed twice with PBS, resuspended in ice-cold 250 mM sucrose/10 mM Tris–HCl (pH 7.4), and homogenized with ten passes of a pre-chilled, zero-clearance homogenizer (Kimble/Kontes). A post-nuclear supernatant was obtained by centrifugation of the samples twice for 10 min at 600g. The heavy membrane fraction was pelleted by centrifugation for 10 min at 10,000g and washed once in the same buffer. The resuspended membranes were loaded on a 1.7 M and 1 M sucrose bilayer and centrifuged in an ultra-centrifuge at 75,000g for 30 min. The layer between the 1.7 M and 1 M sucrose layers was collected and washed in the 250 mM sucrose/10 mM Tris–HCl buffer. The protein concentration was determined by Bradford assay. 40 μg of mitochondria were lysed in 200 μL of the chloroform/isopropanol/NP-40 solution and centrifuged for 10 min at 15,000g.

Supernatants were processed as mentioned above. 20 μL were used per reaction.

### Immunofluorescence

For immunofluorescence analysis cells were fixed in 4% formaldehyde in PBS at 37°C for 20 min, then washed three times with PBS, followed by permeabilization in 0.1% Triton X-100 in PBS and three washes in PBS. The cells were then blocked with 4% fetal bovine serum (FBS) in PBS, followed by incubation with primary antibodies in 4% FBS in PBS, for 1 h at RT. After three washes with PBS, cells were incubated with DAPI and the appropriate anti-species secondary antibodies coupled to Alexa fluorochromes (Invitrogen) (1:3000) for confocal microscopy. After three washes in PBS, coverslips were mounted onto slides using fluorescence mounting medium (Dako).

### Confocal imaging microscopy

For confocal microscopy, stained cells were imaged using a 100× objective lenses (NA1.4) on an Olympus IX81 inverted microscope with appropriate lasers using an Andor/Yokogawa spinning disk system (CSU-X), with a sCMOS camera. A laser power of 10 (for i.e. anti-PRDX3) with a dwell time of 100 μs was used.

For live cell imaging analysis, cells were incubated with MitoTrackerTM Deep Red FM (ThermoFisher) and LysoTrackerTM Red (ThermoFisher) for 15 minutes prior to imaging and washed three times. Time-lapse videos were acquired over the course of 1 min with each channel captured every 0.5 second.

Mitochondrial network morphology was classified using Fiji plugin MitoMAPR (Zhang et al. 2019) which provides “number of networks” and “network length”. At least 10 cells per condition were analyzed in 3 independent experiments.

The fluorescent cholesterol probe was tracked by incubating the cells with a final concentration of 20 μg/ml of TopFluor® Cholesterol (810255, Avanti) and 200 μg/ml of methyl-beta-cyclodextrin (C4555-1G, ThermoFisher) in PBS for 2 min at room temperature. Cells were washed three times and incubated in normal medium for the indicated time. The cells were fixed with 4% PFA and stained with PRDX3 to visualize mitochondria.

### RNA extraction and RNA sequencing

To capture the transcriptome, total RNA from control and SLC25A46 knock-out fibroblast and HeLa cell lines were extracted and purified by using the RNeasy Mini kit (QIAGEN). Three independent biological replicates were collected. RNA quality was tested on a 1% agarose gel and subsequently mRNA sequencing was performed via polyA selection on an Illumina HiSeq 4000 instrument by GENEWIZ, LLC. (South Plainfield, NJ, USA). RNA samples received were measured using Qubit 2.0 Fluorometer (Life Technologies, Carlsbad, CA, USA), and RNA integrity was obtained by 4200 TapeStation (Agilent Technologies, Palo Alto, CA, USA). RNA sequencing library preparation used the NEBNext Ultra RNA Library Prep Kit for Illumina followed by manufacturer’s instructions (NEB, Ipswich, MA, USA). Briefly, mRNA was first enriched with Oligod(T) beads. Enriched mRNAs were fragmented for 15 minutes at 94°C. First strand and second strand cDNA were subsequently synthesized. cDNA fragments were end repaired and adenylated at 3’ends, and universal adapters were ligated to cDNA fragments, followed by index addition and library enrichment by PCR with limited cycles. The sequencing library was validated on the Agilent TapeStation (Agilent Technologies, Palo Alto, CA, USA) and quantified by using Qubit 2.0 Fluorometer as well as by quantitative PCR (KAPA Biosystems, Wilmington, MA, USA). The sequencing libraries were clustered on a single lane of a flow cell. After clustering, the flow cell was loaded on the Illumina HiSeq instrument according to manufacturer’s instructions. The samples were sequenced using a 2×150 Paired End (PE) configuration. Image analysis and base calling were obtained by the HiSeq Control Software (HCS). Raw sequence data (.bcl files) determined from Illumina HiSeq was converted into fastq files and de-multiplexed using Illumina’s bcl2fastq 2.17 software. One mismatch was acceptable for index sequence identification.

### RNAseq preprocessing

Adapters were removed from raw demultiplexed RNAseq reads using bbduk.sh from the BBTools software suite (sourceforge.net/projects/bbmap/) with the settings k = 13 ktrim = n useshortkmers = t mink = 5 qtrim = t trimq = 10 minlength = 25. Next, ribosomal RNA was removed using the same tool, with default settings. HISAT2 (Kim et al. 2019) with settings --no-mixed --no-discordant was used to align the processed reads to hg38. Uniquely mapping reads were quantified using the featureCounts function from the RSubreads package (Liao, Smyth, and Shi 2014) with refSeq gene definitions and with settings isPairedEnd = TRUE useMetaFeatures = TRUE requireBothEndsMapped = TRUE countMultiMappingReads = FALSE allowMultiOverlap = FALSE. The reproducibility of data from all replicates and genotypes was assessed using Principle Component Analysis with genes within the highest quartile of standard deviation across samples.

### Differential gene expression analysis

Analysis of differentially expressed genes in fibroblasts and HeLa cells with SLC25A46 knock-out was carried out using the R/Bioconductor package DESeq2 (version 1.32.0) (Love, Huber, and Anders 2014). Genes with fewer than 10 counts in total across all samples were filtered out prior to all analyses. For pairwise comparisons (KO vs. CTR) in both cell lines, DESeq2 models were run with the design “expression ∼ genotype”, and significantly differential genes were selected based on the thresholds Log2(FC) > |0.3| and adjusted p-value< 0.1. Further analyses were performed with the subset of genes that were significantly differentially expressed upon SLC25A46 knock-out in both fibroblast and HeLa cells (referred to as the SLC25A46-responsive subset).

To identify differences in gene expression under starvation conditions (HBSS treatment) that depended on SLC25A46, all knock-out and control fibroblasts samples with and without HBSS treatment were examined in the same DESeq2 model with the design “expression ∼ genotype + condition + genotype x condition”. Genes with an adjusted p-value < 0.1 and Log2(FC) > |0.3| for the interaction term “genotype x condition” were considered to have significantly different expression under starvation conditions that depended on SLC25A46.

### Gene ontology enrichment analysis

Gene ontology enrichment analysis was performed using the Cytoscape (v.3.8.2.) plug-in ClueGO (v.2.5.8) (Bindea et al. 2009), with genes identified by DESeq2 analysis as being significantly differentially expressed, or with starvation-induced differential expression specifically dependent on SLC25A46. The background for each cell line was defined as all expressed genes. GO term enrichments were selected based on the following criteria: *p*-value cutoff = 0.05, Correction Method Used = Benjamini-Hochberg, Statistical Test Used = Enrichment (Right-sided hypergeometric test), Kappa= 0.4, Min.Percentage = 4, Min GO Level = 3, Max GO Level = 8, Number of Genes = 5, GO Fusion = true, GO Group = true, Over View Term = SmallestPValue, Group By Kapp Statistics = true, Initial Group Size = 1, Sharing Group Percentage = 50.0, Ontology Used = GO_BiologicalProcess-EBI-UniProt-GOA-ACAP, Evidence codes used = All, Identifiers used = SymbolID. Genes that were significantly differentially expressed (Log2(FC) > |0.3| and adjusted p-value < 0.1) and belonging to specific ontology terms were visualized across different samples using clustered heatmaps.

### Co-immunoprecipitation

For cross-link immunoprecipitation, mitochondria enriched heavy membrane fractions (200 μg) from control fibroblasts were chemically cross-linked with 1 mM dithiobis-sulfosuccinimidyl propionate (DSP) (Sigma) in HIM buffer, for 2 h on ice. The reaction was stopped by adding glycine pH 8.0 at 70 mM final concentration for 10 min on ice. Mitochondria were pelleted, rinsed once, and extracted in 200 μl of lysis buffer (10 mM Tris–HCl pH 7.5, 150 mM NaCl, 1% n-dodecyl-D-maltoside (DDM) (Sigma), and complete protease inhibitors (Roche)) on ice for 30 min. The extract was centrifuged at 20,000g at 4°C for 20 min, and the supernatant was pre-cleared overnight with non-coated Dynabeads Protein A (Invitrogen) to reduce non-specific protein binding to the beads. Binding of indicated antibodies to Dynabeads Protein A (Invitrogen) was performed overnight. Antibodies were then cross-linked to the beads using 20 mM dimethyl pimelimidate (DMP) (Sigma). The immunoprecipitation reaction was performed overnight at 4°C. Beads were washed with lysis buffer and samples were eluted using 0.1 M glycine pH 2.5/0.5% DDM, precipitated with trichloroacetic acid and analyzed by mass spectrometry at the IRCM (Institut de Recherches cliniques de Montreal).

### Antibodies

Antibodies directed against the following proteins were used in this study: SLC25A46 (Santa Cruz; G2-SC-515823 and Proteintech; 12277-1-AP), MFN2 (Cell Signaling; 11925S), MFN1 (Cell Signaling; 14739S), OPA1 (BD bioscience, 612607) for WB, VDAC1 (Abcam; ab14734), 4E-BP (Cell Signaling; 9644S), Cav1 (BD bioscience; 610060). HMGCR (Invitrogen; MA5- 31335) PRDX3 (in house), ATP5A1 (Abcam; ab14748); the secondary antibodies: anti-rabbit Alexa Fluor 488 (Invitrogen; A-21206), anti-mouse Alexa Fluor 647 (Invitrogen; A-31571), anti-rat Alexa Fluor 594 (Invitrogen; A-11007),

### Statistical analysis

All data are reported as means ± SEM or means ± SD as indicated in the figure legend. Statistical significance was determined using the indicated tests. P-values < 0.05 were considered statistically significant and labeled as follows: *P < 0.05, **P < 0.01, ***P < 0.001 and ****P<0.0001.

## ACKNOWLEDGEMENTS

This research was supported by a grant from the CIHR to EAS (FRN178373). We thank Ivan Topisirovic for valuable advice.

## AUTHOR CONTRIBUTIONS

JS led the project, designed and performed the experiments and co-wrote the manuscript. KW analysed the trancriptomic data and co-wrote the manuscipt. AJ contributed to the experimental design. HA designed and performed the BioID experiments and co-wrote the manuscript. OL supervised the transcriptomics analysis. EAS helped with experimental design, supervised the project, and co-wrote the manuscript.

---

Figure S1: SLC25A46 knock-out cells have an altered mRNA expression. Transcriptomics data was analysed for control and SLC25A46 knock-out cell lines in fibroblasts and HeLa cells. (a) Principle component analysis of gene expression quantified by RNA-seq in control and SLC25A46 knock-out HeLa cells. (b) Quantification of SDS-PAGE of control fibroblasts and the SLC25A46 knock-out cell line not treated or supplemented with 20 μM of cholesterol (Figure 6d). VDAC was used as a loading control. P-values were calculated using a two-tailed, unpaired t-test, n=3 independent experiments. No significant differences were found. (c) Principle component analysis of gene expression quantified by RNA-seq in control and SLC25A46 knock-out fibroblasts upon no treatment (unstarved) and treatment with HBSS for 4 hours (starved).

**Figure S2: Cholesterol trafficking is dependent on SLC25A46.** Immunofluorescent analysis of control human fibroblasts and SLC25A46 knock-out cells. (a) Calnexin was used as an ER marker (in magenta). PRDX3 was used as a mitochondrial marker (in green). (b) Cholesterol trafficking using LysoTracker® (in green) as a lysosomal marker and TopFluor® (in magenta). Cells were pulsed for 2 min with the fluorescent cholesterol probe. Scale bars: 10 μm. (c, d) Heatmap of up- and downregulated SLC25A46-responsive genes in control and SLC25A46 knock-out fibroblasts belonging to a GO term: proteins expressed in vesicles (GO:00031982) (c) and lysosomes (GO:0005764) (d). Log2 fold changes are centered and represented as z-scores.

Figure S3: Starvation induces upregulation in transcripts of NF-κB signalling, response to ER-stress, MAPK-cascade and JNK cascade in the SLC25A46 knock-out cell line. Heatmap of SLC25A46-dependent differentially expressed genes in control and SLC25A46 knock-out fibroblasts in normal condition (unstarved) and in HBSS treated condition (starved) for transcripts in NF-kappaB signalling (GO:0007249) (a), response to endoplasmic reticulum stress (GO:0034976) (b), MAPK cascade (GO:0000165) (c), and Regulation on JNK cascade (GO:0046328) (d). Genes displayed are those where HBSS-induced changes in expression were dependent on SLC25A46 in the DESeq2 model with the design “expression ∼ genotype + condition + genotype x condition” (adjusted p-value < 0.1 and Log2FC > |0.3| for the “genotype x condition” term). Log2 fold changes are centered and represented as z-scores..

