## Supplementary figures and images for "SLC25A46 is in contact with lysosomes and plays a role in mitochondrial cholesterol homeostasis"

### Supplementary Fig. 1

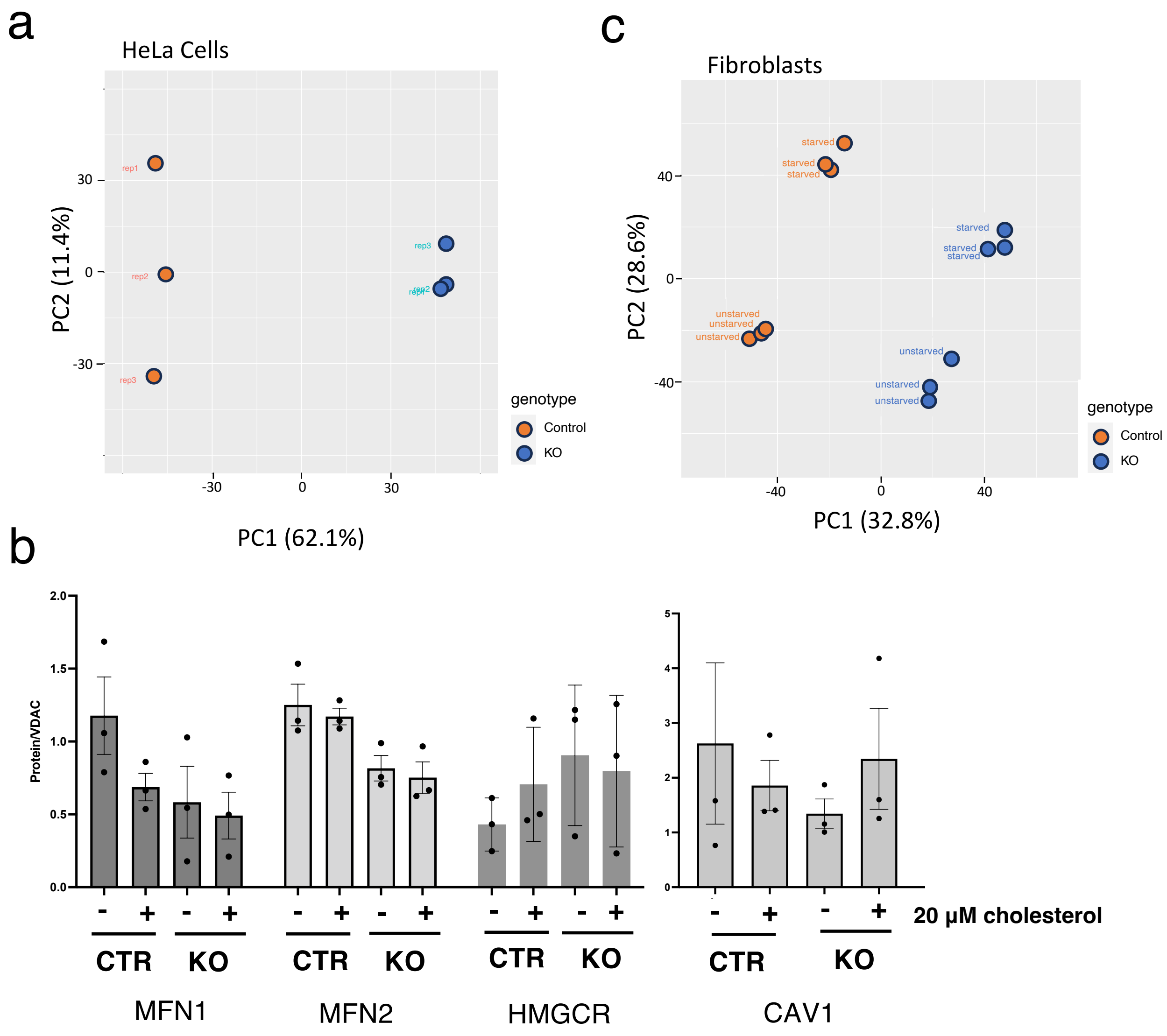

### Supplementary Fig. 2

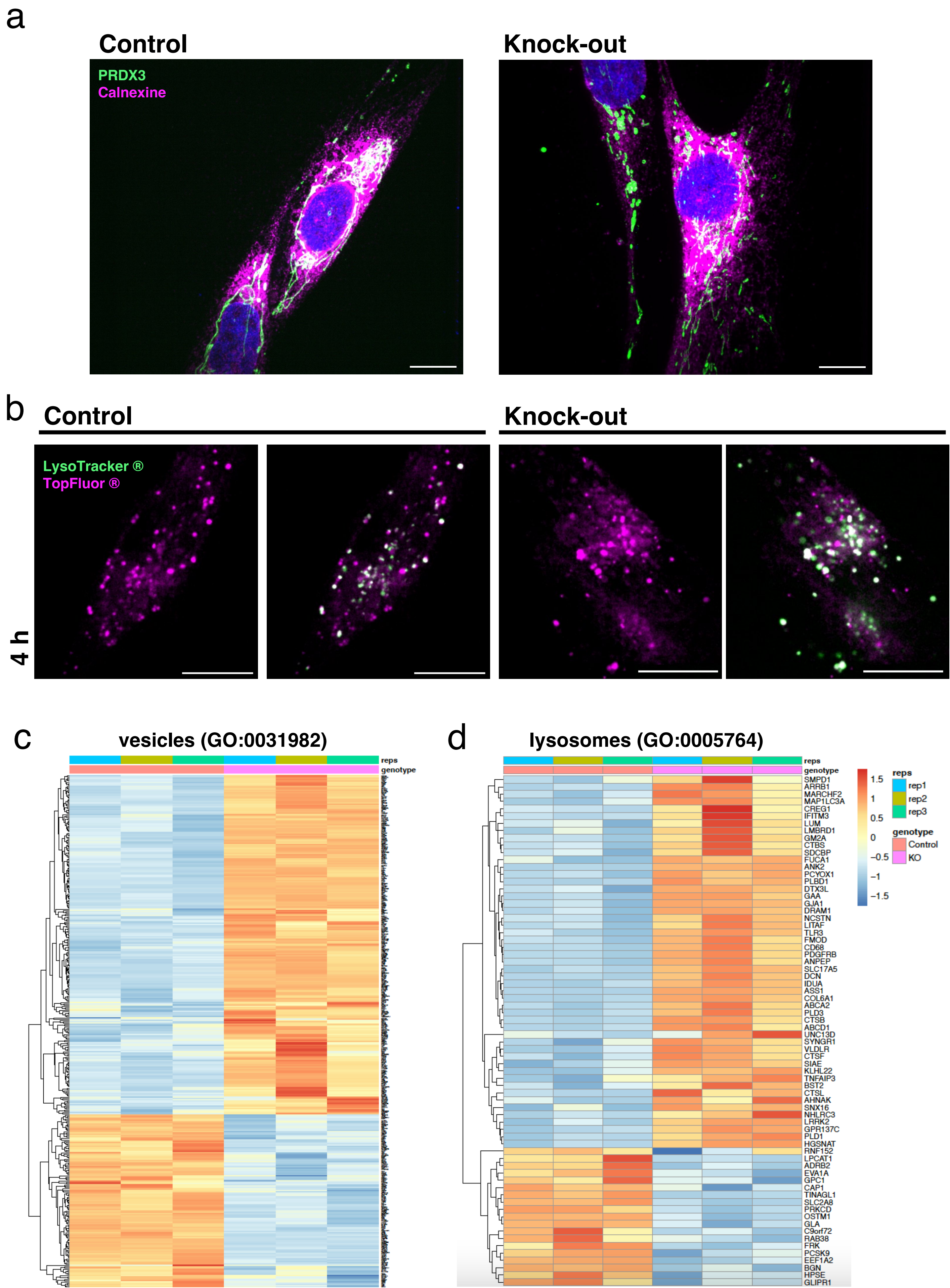

### Supplementary Fig. 3ab

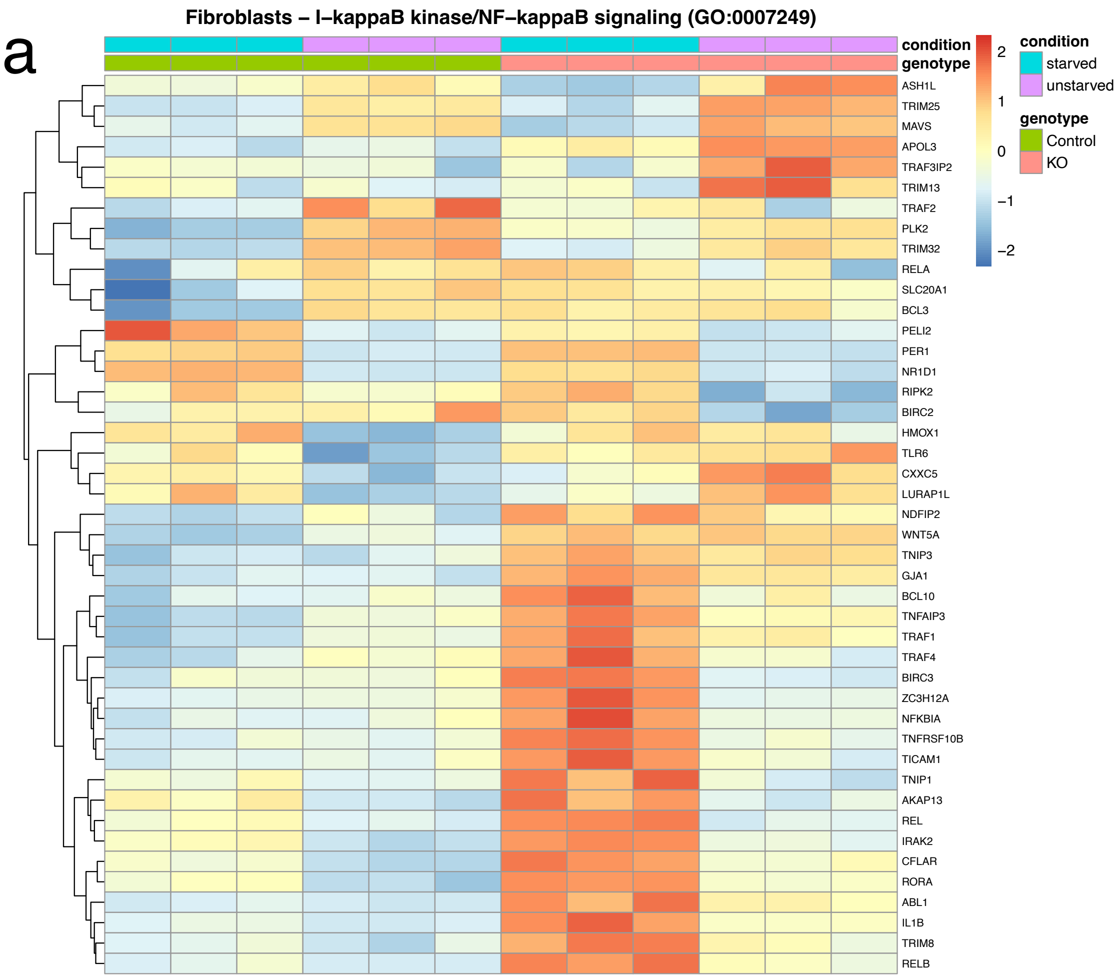

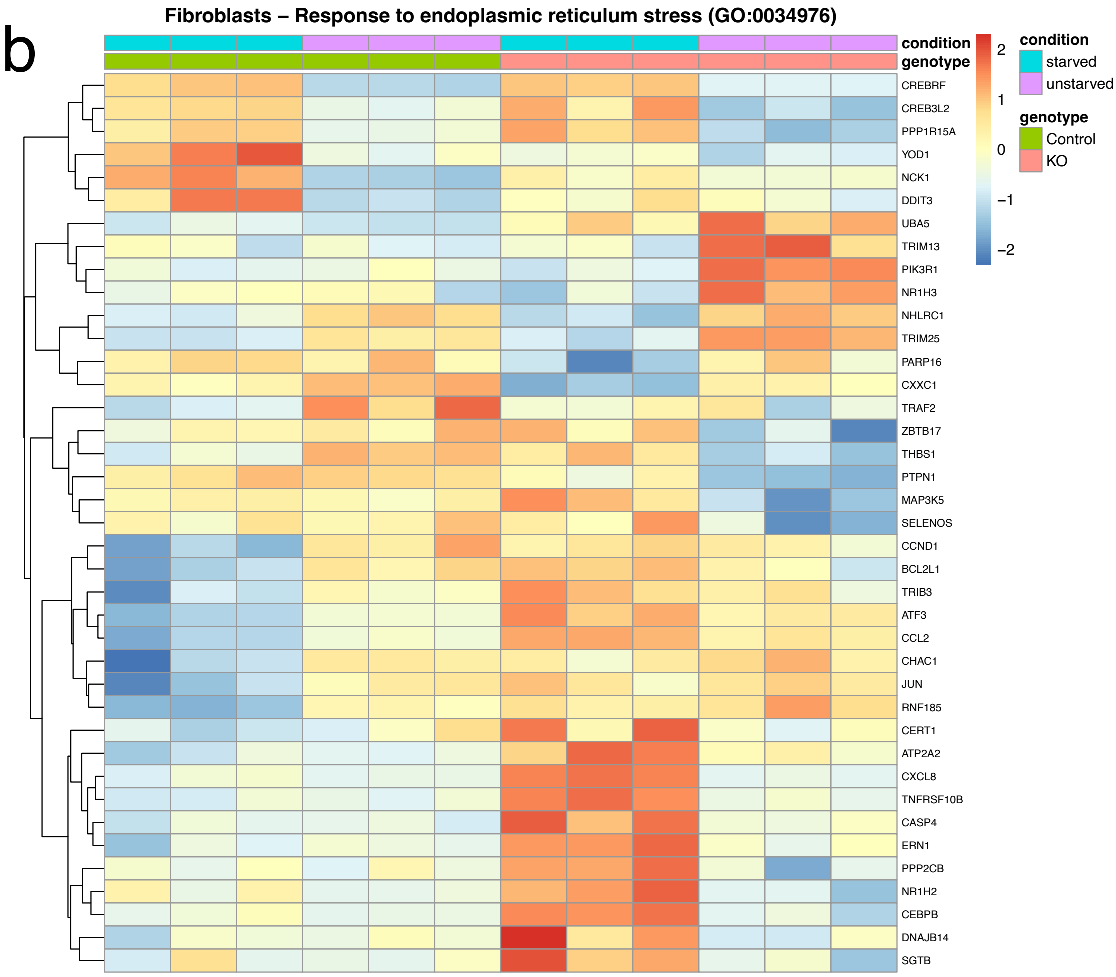

### Supplementary Fig. 3cd

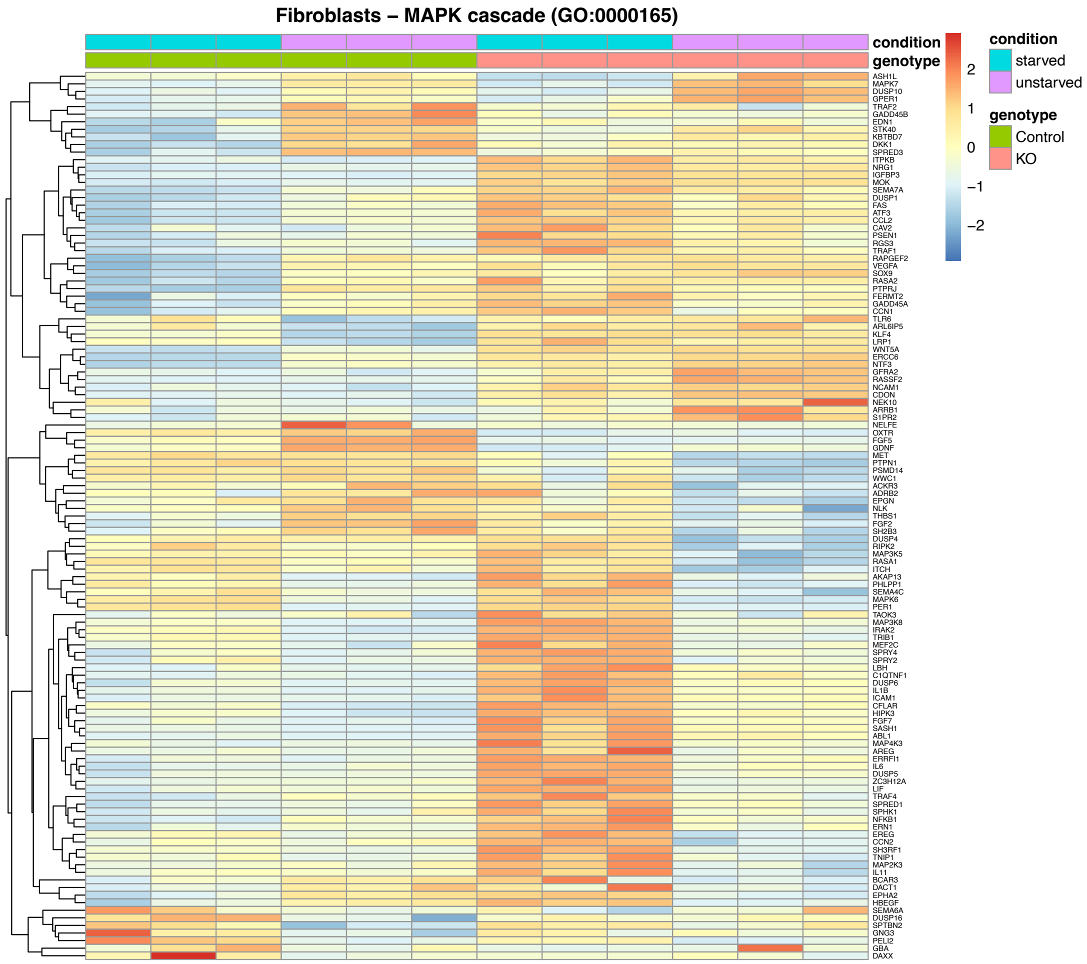

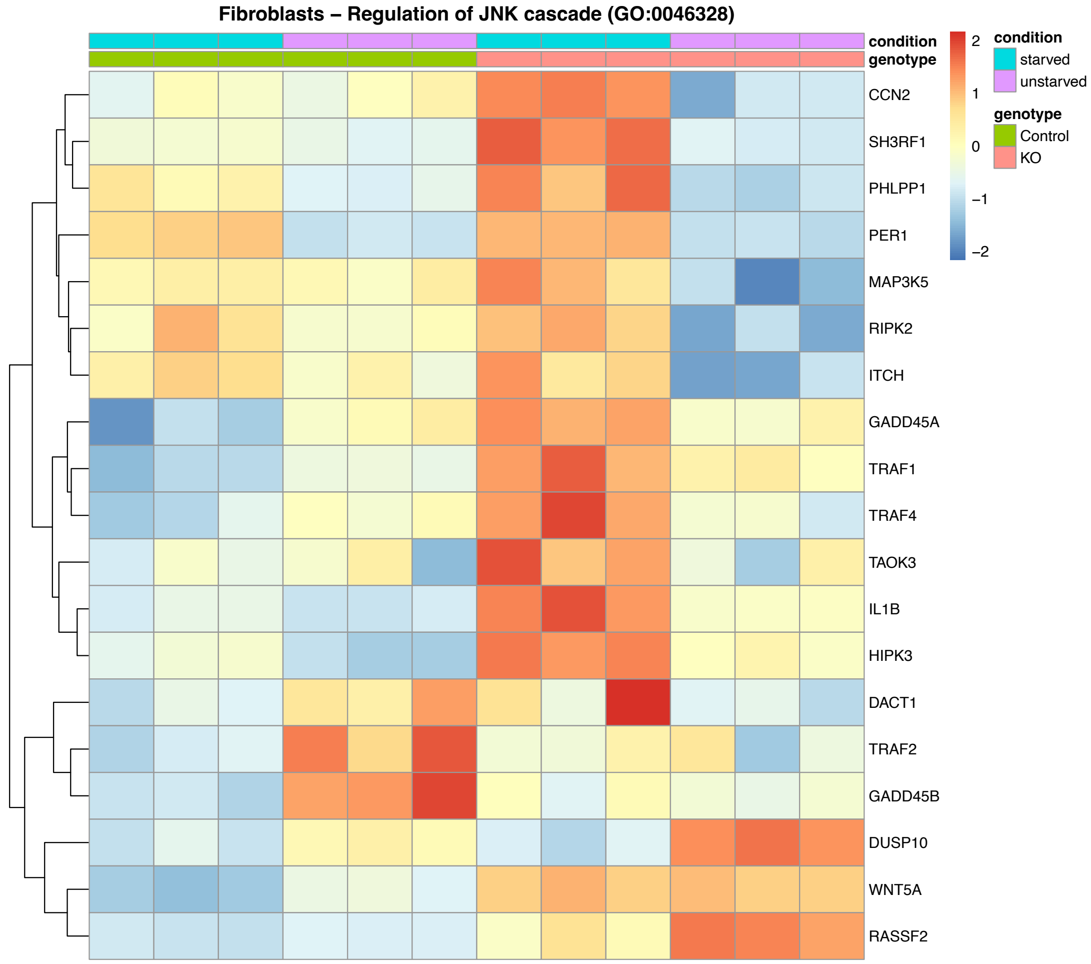
